# Altered Expression of microRNAs Implicated in Hematopoietic Dysfunction in the Extracellular Vesicles of Bone Marrow-Mesenchymal Stromal Cells in Aplastic Anemia

**DOI:** 10.1101/2024.04.20.590382

**Authors:** Jyotika Srivastava, Kavita Kundal, Bhuvnesh Rai, Pragati Saxena, Naresh Tripathy, Sanjeev Yadav, Ruchi Gupta, Rahul Kumar, Soniya Nityanand, Chandra Prakash Chaturvedi

## Abstract

Recently, we have reported that extracellular vesicles (EVs) from the bone marrow mesenchymal stromal cells (BM-MSC) of aplastic anemia (AA) patients inhibit hematopoietic stem and progenitor cell (HSPC) proliferative and colony-forming ability and promote apoptosis. One mechanism by which AA BM-MSC EVs might contribute to these altered HSPC functions is through microRNAs (miRNAs) encapsulated in EVs. However, little is known about the role of BM-MSC EVs derived miRNAs in regulating HSPC functions in AA. Therefore, we performed miRNA profiling of EVs from BM-MSC of AA (n=6) and normal controls (NC) (n=6), to identify differentially expressed miRNAs carried in AA BM-MSC EVs. DEseq2 analysis identified 34 significantly altered mature miRNAs in AA BM-MSC EVs. Analysis of transcriptome dataset of AA HSPC genes identified that 235 differentially expressed HSPC genes were targeted by these 34 EV miRNAs. The pathway enrichment analysis of 235 HSPC genes revealed their involvement in pathways associated with cell cycle, proliferation, apoptosis, and hematopoiesis regulation, thus highlighting that AA BM-MSC EV miRNAs could potentially contribute to impaired HSPC functions in AA.

## 1. Introduction

Idiopathic acquired aplastic anemia (AA) is a state of bone marrow failure characterized by a fatty hypoplastic marrow and multilineage cytopenia’s ^1^. Several studies have reported that perturbations in the bone marrow (BM) microenvironment in AA may suppress hematopoietic stem and progenitor cells (HSPC)^2–4^. The BM mesenchymal stromal cells (BM-MSC) are a key cell of the BM microenvironment and have immunomodulatory and hematopoietic supporting functions^5,6^.

Data from our and other labs have confirmed that BM-MSC of AA patients have cellular and molecular defects, implying a possible role of these defects being responsible for their reduced ability in sustaining HSPC functions when compared to their normal counterparts^4,7–12^. However, the mechanism by which BM-MSC impart their functional effect on HSPC in AA remains elusive. Recent evidence suggests that MSC affect the functions of HSPC via paracrine release of certain hematopoietic cytokines, growth factors and extracellular vesicles (EVs)^13–16^.

Extracellular vesicles (EVs) are 20-1000 nm sized nano-vesicles, which serve as the key mediators of intercellular communication by modulating the biological functions of their target cells^17,18^. Several studies have reported that BM-MSC derived EVs can affect self-renewal, proliferation, and differentiation of HSPC in normal physiological conditions analogous to that of BM-MSC^19–21^. Moreover, in hematological disorders such as acute myeloid leukaemia (AML), myelodysplastic syndrome (MDS) and multiple myeloma (MM), it has been reported that BM-MSC derived EVs can impair the hematopoiesis supportive functions of HSPC^22–24^.

We have recently reported that BM-MSC EVs from AA patients can inhibit the proliferative and colony-forming ability of HSPC by enhancing apoptosis, thus highlighting the altered hematopoiesis supporting properties of AA BM-MSC EVs^25^. Mechanistically, EVs can profoundly alter the target cell phenotype by altering gene expression and associated pathways by releasing bioactive factors such as miRNAs encapsulated within them^17^. These miRNAs can orchestrate the function of HPSC via post-transcriptional regulatory mechanisms^26–28^. Thus, suggesting that alterations in the expression of these EV miRNAs can hamper the HPSC functions. Supporting this hypothesis, a recent study by Guidice et al., 2018 demonstrated that few plasma exosomal miRNAs have a role in the dysregulated expansion and cell survival of HSPC in AA patients ^29^.

However, the role of miRNAs derived from the EVs of AA BM-MSC in HSPC dysfunction remain obscure to date. Thus, in the present study we have performed global miRNAs profiling of EVs from the BM-MSC of AA patients and controls to identify significantly altered miRNAs. Furthermore, utilizing the transcriptome dataset of the HSPC of AA and controls available in the NCBI repository, we aim to identify HSPC genes which are targeted by these differentially expressed EV miRNAs. Additionally, in-silico functional analysis of target HSPC genes was carried out to identify regulatory pathways and hub genes which could be potentially involved in impeding HSPC functions. Overall, the study underlines the role of AA BM-MSC EVs-derived miRNAs in the disease pathobiology.

## 2. Results

### 2.1. EV from BM-MSC of AA patients and normal control (NC) exhibit similar size, morphology, and expression of EVs specific markers

The BM-MSC of AA patients and NC exhibited the presence of MSC markers; CD73, CD90 and CD105 and had negligible expression of hematopoietic markers; CD34, CD45 and HLA-DR. The evaluation of differentiation potential of BM-MSC revealed that AA BM-MSC had a higher adipogenic differentiation potential and a reduced osteogenic and chondrogenic differentiation potential as reported previously^4,11^ (see **Supplementary Fig. S1**). EVs from BM-MSC of AA patients and NC had comparable particle size (145.3nm vs 146.9nm, *p-value-0.87*), and concentration (6.55e+007 vs 5.22e+007*, p-value-0.48*) (**Fig. 1a-c**). Both AA and NC EVs had cup shaped, circular morphology as analyzed by TEM (**Fig. 1d**) and exhibited the expression of CD63, an EV marker (87.43 vs 83.07, *p-value-0.43*) as confirmed by flow-cytometry (**Fig. 1e-f**). Aligning with Minimal information for studies of extracellular vesicles (MISEV 2023) guidelines^30^, the EVs from both groups exhibited the presence of CD63, CD81, TSG101 and had negligible expression of calnexin (CNX) (a negative EV marker, ER associated marker) as confirmed by western blotting (**Fig. 1g**). The size estimation by TEM and original western blots images of AA and NC BM-MSC EVs is provided in **Supplementary Fig. S2.**

**Figure 1:**
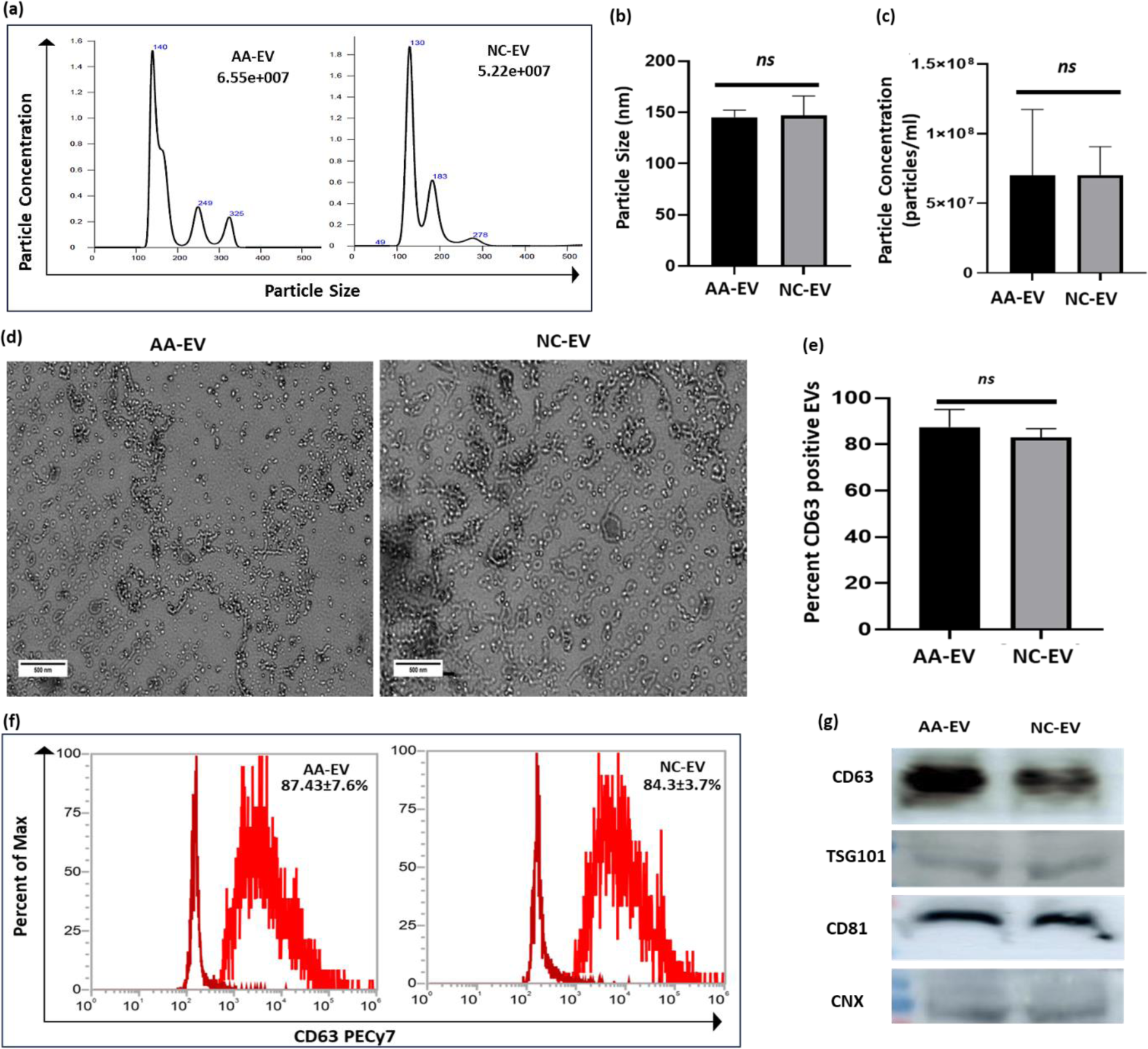
Characterization of EVs from BM-MSC of AA and NC. **(a)** Representative histogram showing particle size and concentration of EVs from AA and NC BM-MSC. **(b-c)** Bar graphs showing comparison of particle size and concentration of EVs from AA and NC BM-MSC, **(d)** Representative photomicrographs showing EVs morphology as analyzed by TEM, **(e)** Bar graph and **(f)** representative histograms comparable expression of CD63 in the EVs from AA and NC BM-MSC, **(g)** Immuno-blot images exhibiting the presence of CD63, TSG101 and CD81 and negligible expression of CNX. **Note:** TEM: Transmission electron microscopy; CD: cluster of differentiation; TSG: Tumour susceptibility gene, CNX: Calnexin; ns-non-significant.

### 2.2. Differential expression of miRNA in the EVs derived from AA BM-MSC

Global miRNA profiling was performed to EVs from BM-MSC of AA patients and NC identify the differential expression of precursor and mature miRNAs. The normalization of the read count was performed on 6 AA and 6 NC samples. Out 6 AA samples we had to exclude two samples because of the poor read quality (**Fig. 2a**). We found that out of identified 702 miRNAs, 57 miRNAs were significantly altered in AA BM-MSC EVs compared to NC BM-MSC EVs (**Fig. 2b**). The heatmap of the differentially expressed miRNAs was generated using ComplexHeatmap tool embedded in R (**Fig. 2c**). Out of 57 differentially expressed miRNAs, 23 were precursor and 34 mature miRNAs. We further proceeded with the mature miRNAs, out of 34 miRNAs, 9 were upregulated and 25 were downregulated. The tabulated form of differentially expressed miRNAs, both precursor and mature with fold change and *p-value* is given it the **Supplementary Table S1 and Table S2**.

**Figure 2:**
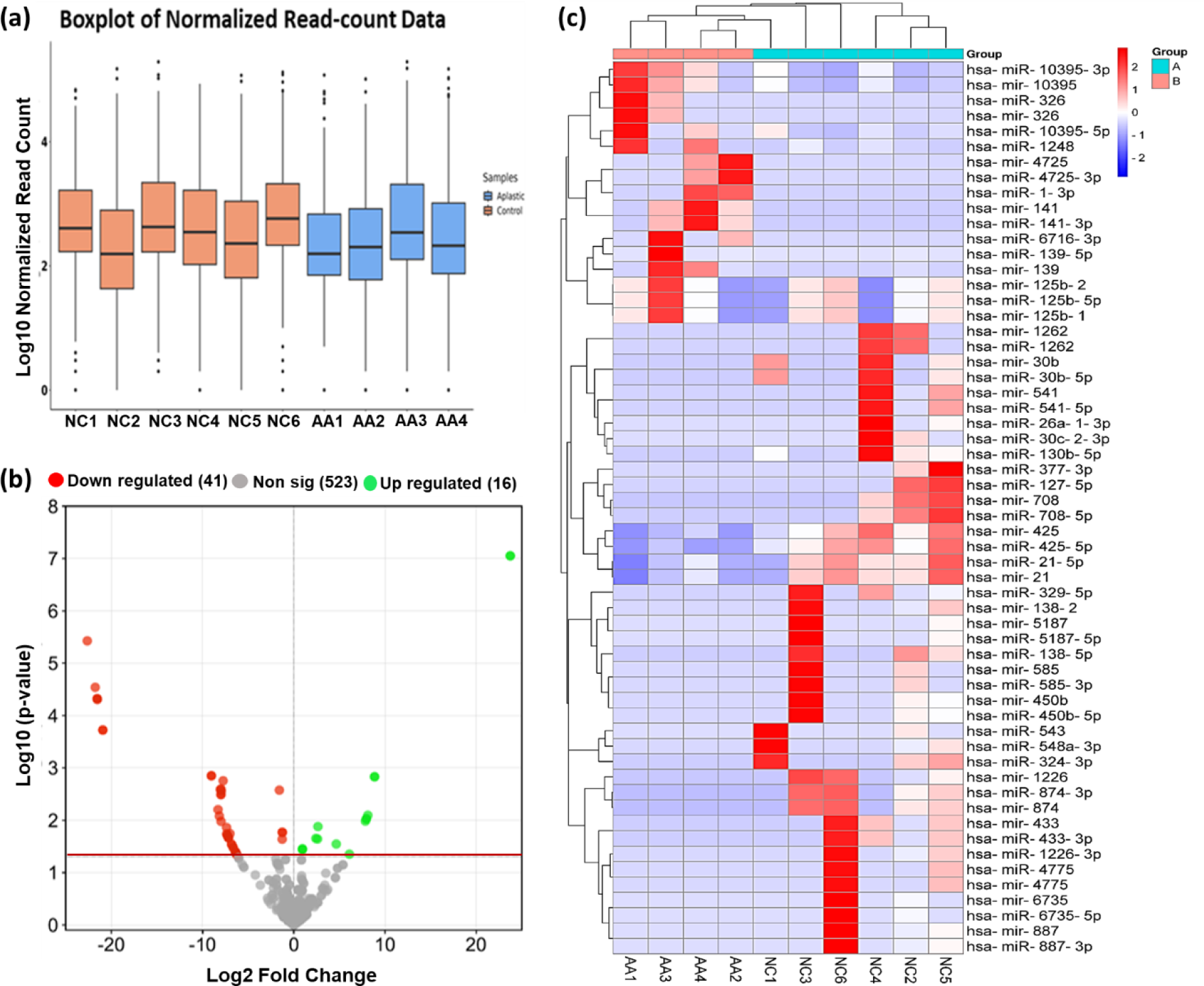
Differential expression of miRNAs in EV of AA BM-MSC in comparison to NC BM-MSC. **(a)** Log expression values of normalized read counts across all samples used for miRNA analysis of EVs from AA BM-MSC (n=4) and NC BM-MSC (n = 6), **(b)** Volcano plot of differentially expressed EV miRNAs in AA BM-MSC when compared to NC BM-MSC where green dots represent upregulated miRNAs (log2FC > 1.0) and red dots represents downregulated miRNAs (log2FC < -1.0) and having *p-value <0.05*, **(c)** Heatmap showing hierarchical clustering of differentially expressed miRNAs in the EVs from BM-MSC of AA (n=4) and NC (n = 6) with *p-value <0.05*. The color key indicates Z score calculated from the TPM values; dark blue = lowest and, dark red = highest.

### 2.3. Identification of pathways associated with differentially expressed EV miRNAs

Further we performed in-silico functional analysis of differentially expressed miRNAs in EVs of AA BM-MSC to identify the Gene ontology (GO) pathways and biological processes using miEAA^31^ having *p-value <0.05*. The pathways analysis revealed enrichment of pathways such as, DNA damage responses to ATM, p53 signalling, PI3K-Akt signalling, MAPK signalling, TNF signalling, mTOR signalling etc. which are involved in regulation of cell survival and cell death (**Fig. 3a**). Also, we found that 31 miRNAs were involved in regulation of biological processes of our interest such as cell cycle arrest, negative cellular proliferation, apoptosis, vesicle mediated transport, T-cell differentiation, cellular senescence, and hematopoiesis as represented in the chord plot (**Fig. 3b**). The gene set enrichment analysis (GSEA) of significantly altered miRNAs showed enrichment of two pathways viz., apoptosis and DNA damage responses (**Fig. 3c**).

**Figure 3:**
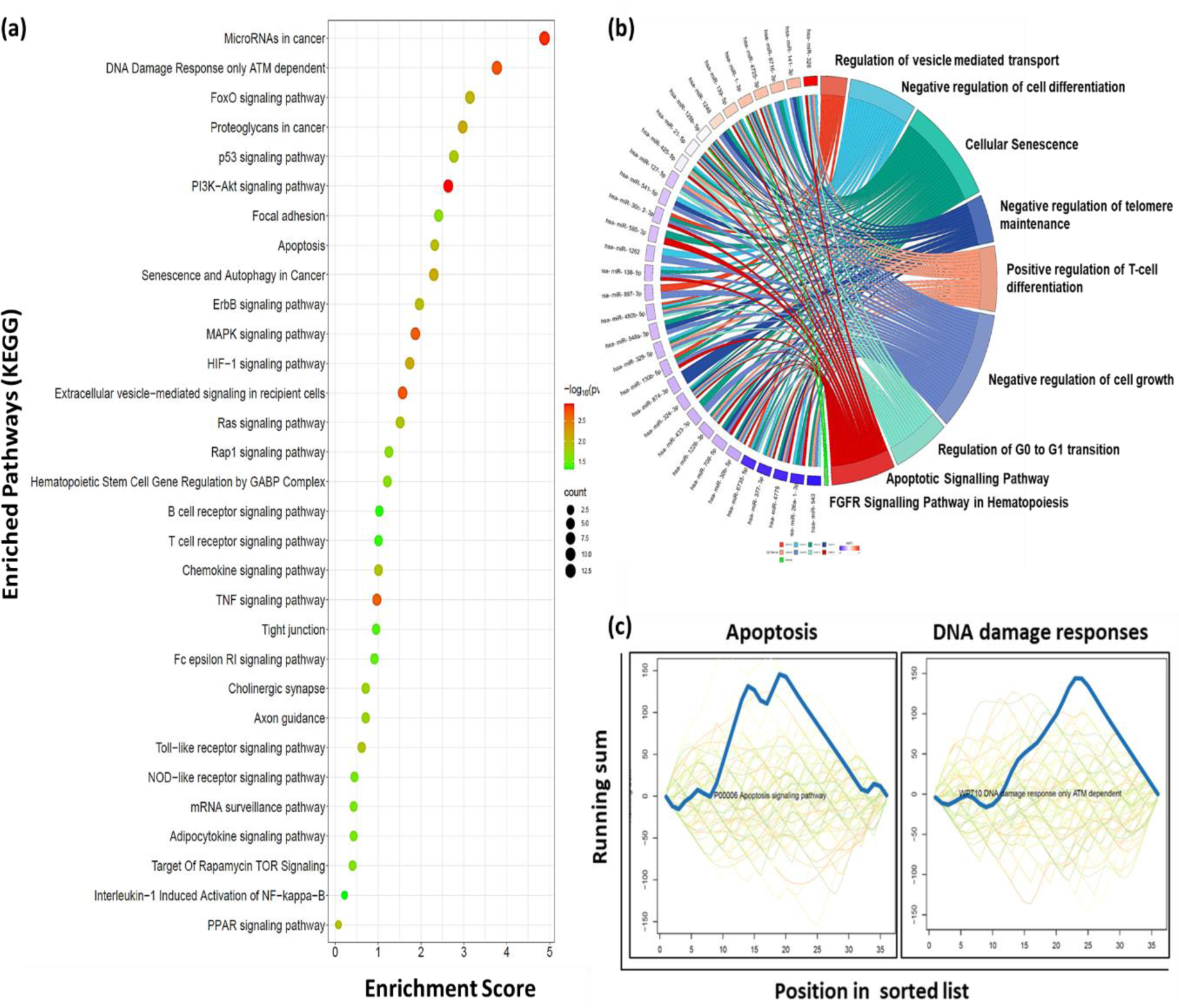
Functional enrichment of differentially expressed miRNAs in EVs of AA BM-MSC. **(a)** Bubble plot showing enrichment of significant pathways targeted by differentially expressed EV miRNAs. The color of the dot represent -log10 p-value and the size of the dot represents numbers of miRNAs involved in regulation of a particular pathway. **(b)** Chord plot showing the miRNAs involved in the regulation of biological processes of our interest and, **(c)** GSEA analysis of significantly altered miRNAs showing enrichment of apoptosis and DNA damage responses pathway. **Note:** DE-differentially expressed; GSEA-Gene set enrichment analysis

### 2.4. EV miRNAs from BM-MSC of AA patients target HSPC genes

Further, we wanted to explore that whether these dysregulated EV miRNAs are involved in the regulation of HSPC genes which are altered in AA patients. For that first we identified that gene targets of 34 differentially expressed mature miRNAs using miRNet 2.0^32^ which predicted a total of 6294 target genes. Next, we analyzed a bulk RNA sequencing dataset of HSPC from AA patients and controls available in the gene expression omnibus (GEO) database having accession number GSE165870. We performed feature counts DESeq2 analysis to identify the differentially expressed HSPCs genes in AA patients in comparison to controls. The volcano plot showed the significantly upregulated (red dots), downregulated (green dots) and non-significant (black dots) genes in HSPCs of AA patients (**Fig. 4a**). Further we identified the 235 overlapping genes between differentially expressed miRNAs target genes and HSPC genes as represented by Venn diagram (**Fig. 4b**). The heatmap showed clearly distinct expression profile of 235 HSPC genes with 166 up and 69 down-regulated genes in AA patients (**Fig. 4c**). Moreover, we found a significant enrichment of several pathways such as PI3K-Akt, mTOR, HIF-1, NF-kappa B signalling, NK cell mediated cytotoxicity, chemokine signalling, autophagy, and apoptosis etc, regulated by the 235 HSPC genes that were targeted by EVs miRNAs of AA BM-MSC. (**Fig. 4d**).

**Figure 4:**
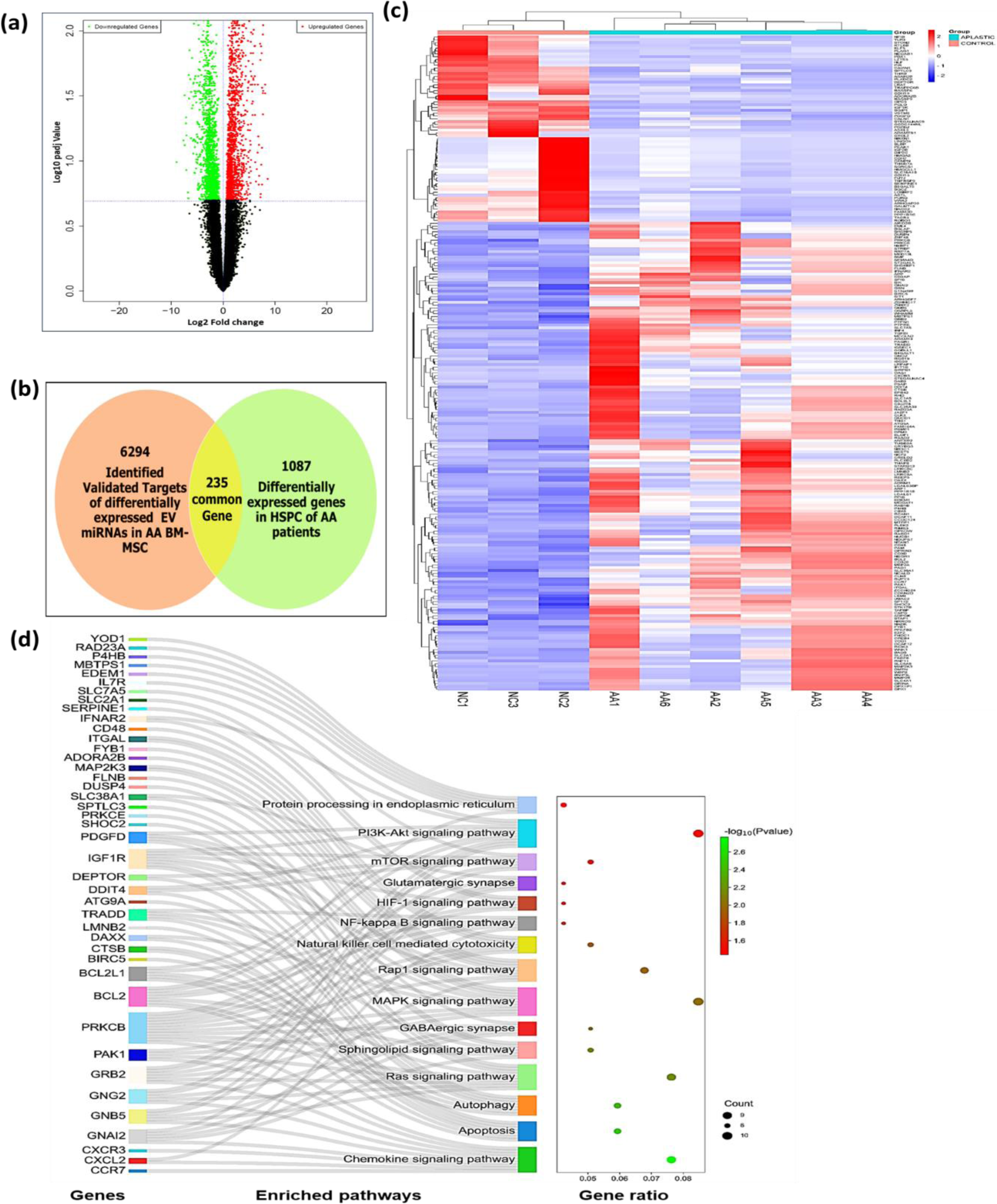
Differential expression of HSPC genes in AA patients in comparison to controls. **(a)** The volcano pot showing the DEGs in the HSPC of AA patients where green dots are significantly downregulated and red dots are significantly upregulated, **(b)** Venn diagram showing 235 common genes between identified 6249 target genes of differentially expressed EV miRNAs and 1087 differentially expressed genes in HSPC of AA patients **(c)** Heatmap representing the up and downregulation of the 235 target HSPC genes and, **(d)** DAVID analysis showing pathways, their gene-ratio, p-value and, HSPC genes involved in regulation of enriched pathways as represented by the Sankey and bubble plot.

### 2.5. miRNA-mRNA network construction and identification of key clusters using Protein-protein (PPI) network analysis

The dysregulated miRNAs and target HSPC gene regulatory network was constructed using miRNet 2.0 (**Supplementary Fig S3**). Further, we submitted 235 common DEGs into the STRING v11 PPI network for identifying significant gene clusters by using the MCODE (molecular complex detection) plugin embedded in the Cytoscape. The analysis identified five significant MCODE clusters of HSPCs genes (**Fig. 5a-b**). The functional enrichment analysis revealed the association of five MCODE clusters with GPCR signalling, chemokine signalling interferon gamma signalling, PID NFAT pathway, apoptosis, autophagy, vesicle mediated endocytosis and glycosphingolipid biosynthesis in a cluster-dependent manner (**Fig. 5c**). Thus, suggesting that these gene clusters could be regulated by the AA BM-MSC EVs encapsulated miRNAs.

**Figure 5.**
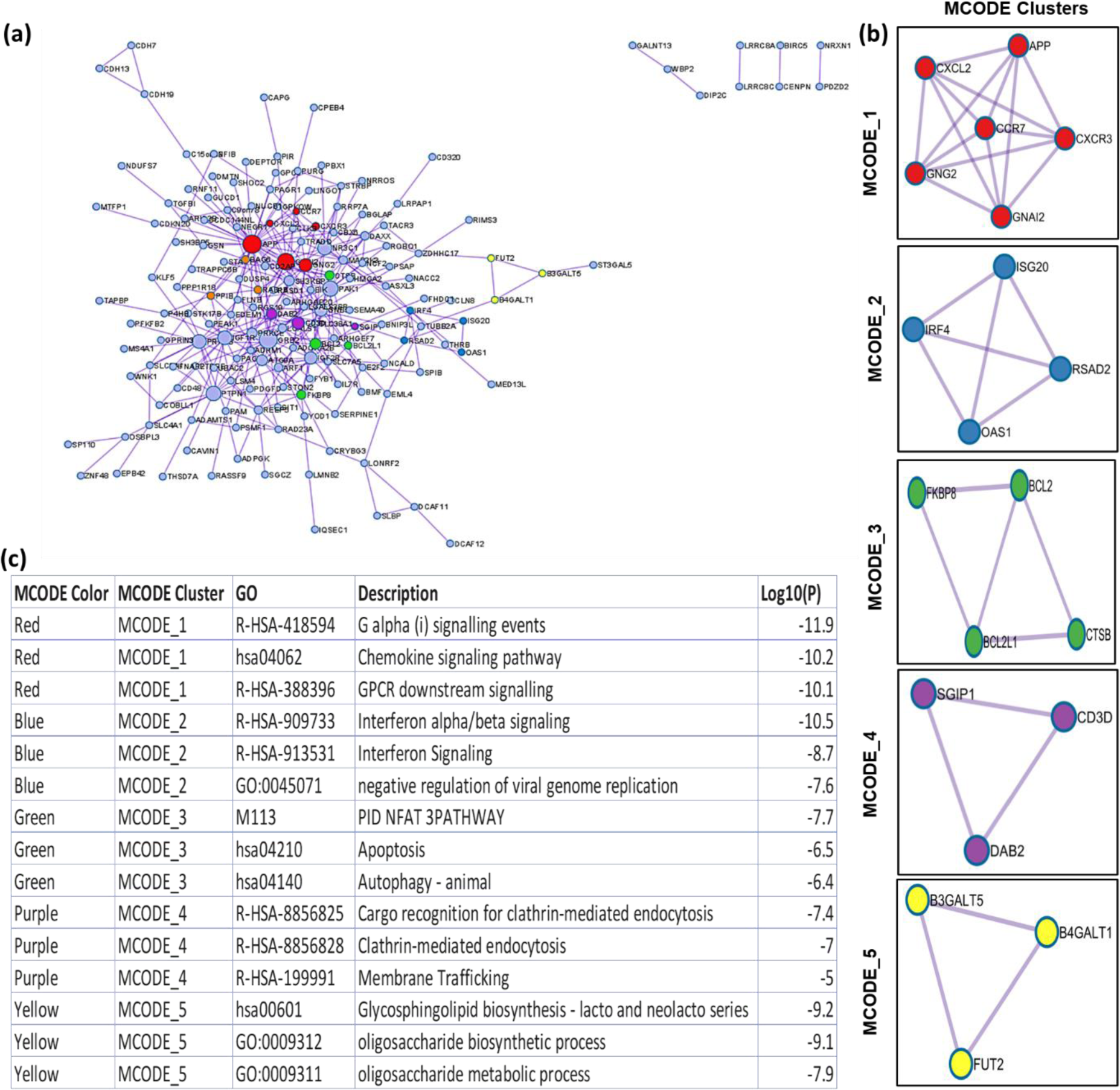
STRING PPI network analysis and MCODE cluster identification and functional analysis. **(a)** PPI network analysis showing different MCODE cluster out of 235 gene**, (b)** The different color coded MCODE clusters showing highly interactive genes in individual clusters and**, (c)** Pathway enriched for different MCODE clusters along with their respective -log10 *p-value*.

### 2.6. Identification of hub genes and their regulatory network

We further screened the top 10 highly interactive genes (hub genes) out of the 235 target genes. The differentially expressed mRNAs were entered as query into the STRING database for a PPI network analysis using CytoHubba embedded in Cytoscape. The top ten hub genes with the highest degree of interactions were identified as: APP (degree = 41), GRB2 (degree = 19), GNAI2 (degree = 18), PAK1 (degree = 16), and PTPN (degree = 15), PRKCB (degree=14), IGF1R (degree=14), NR3C1 (degree=14), GNG2 (degree=13) and, IGF2R (degree=12) (**Fig. 6a**). The APP, GRB2, GNAI2, PAK1, PTPN, PRKCB, NR3C1, and GNG2 were significantly upregulated whereas IGF1R and IGF2R were downregulated in the HSPC of AA patients. The GO analysis of hub genes revealed the involvement of these genes in several biological processes such as signal transduction, insulin receptor signalling pathway, apoptosis, GPCR signalling, cellular ROS etc (**Fig. 6b**). Further, a KEGG pathway analysis of the top 10 hub HSPC genes showed enrichment of B cell receptor signalling, chemokine signalling, EGFR, ErBb, MAPK signalling, NK cell mediated cytotoxicity, Rap1 signalling, PI3k-Akt signalling, mTOR and Ras signalling pathways (**Fig. 6c**). The GO molecular functions and cellular component of the Hub genes is provided in the **Supplementary Fig. S4 and Table S3.**

**Figure 6.**
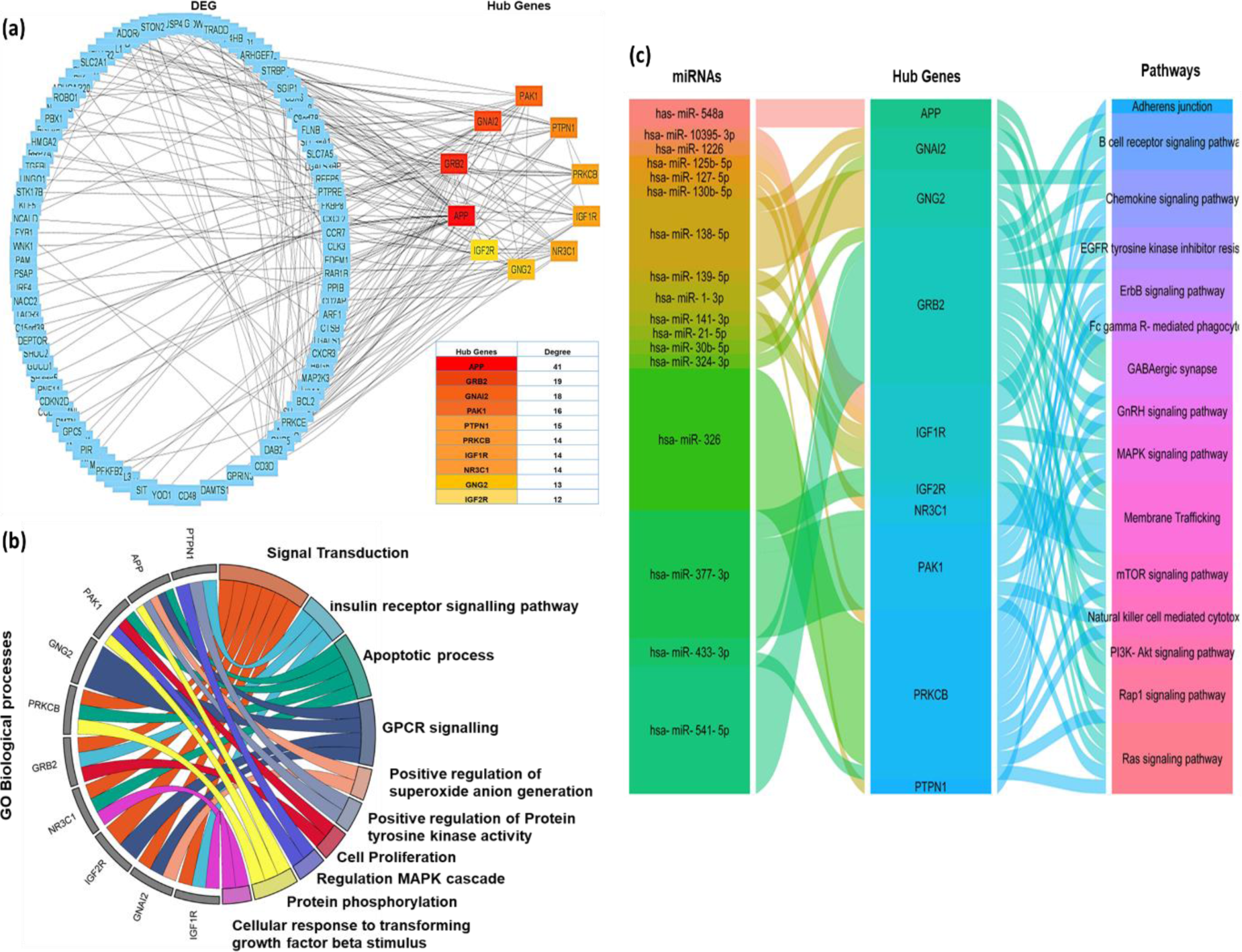
The STRING analysis for hub genes identification and functional analysis. **(a)** PPI network construction and identification of top 10 hub genes where red represent gene with highest degree and yellow with the lowest degree, **(b)** Chord plot representing the regulation of enriched GO biological processes targeted by the Hub genes (color-coded) and, **(c)** Alluvial plot representing the comprehensive network of BM-MSC EV miRNAs involved regulation of hub HSPC genes and associated pathways. Note: GO=Gene ontology; DEG-Differentially expressed genes.

## 3. Discussion

An “empty bone marrow with depletion in the HSPC number” is one of the characteristics of patients with AA. The current understanding of the disease etiopathology includes cytotoxic T-cell activation and abnormal bone marrow microenvironment^4–6,11,12,33^. BM-MSC are the key cells of the BM microenvironment that supports HSPC functions by direct cell contact and paracrine release of growth factors and EVs^5,6,13–16^. We have previously demonstrated that BM-MSC EVs from AA patients hamper the proliferative, colony-forming capacity and enhance apoptosis of HSPC^25^. However, the exact mechanism by which AA BM-MSC EVs exert their functional effect on HSPC is unclear. Filling this gap, the present study revealed that AA BM-MSC-derived EVs have altered expression of miRNAs that could directly impact genes/pathways involved in HSPC cell proliferation, colony-formation capacity, apoptosis, etc., thus highlighting that the EVs of BM-MSC of AA patients mediate impaired HSPC functions via the transfer of encapsulated miRNAs.

BM-MSC plays a vital role in maintaining hematopoietic homeostasis via the release of EVs, which are critical mediators of cellular communication^13,15,16^. However, studies investigating the role of BM-MSC EVs in AA are scarce. It has been shown that BM-MSC EVs from AA patients induced apoptosis and reduced the cellular proliferation and colony forming ability of HSPC^25^. Another study has shown that EVs from healthy BM-MSC potentially ameliorate bone marrow failure symptoms in the murine AA model^36^. These findings imply that BM-MSC EVs significantly regulate HSPC functions under normal or pathological hematopoiesis conditions. Mechanistically, EVs are shown to impart their functional effect via the transfer of miRNA cargoes to HSPC, thereby modulating the target genes and pathways in HSPC regulating normal or malignant hematopoiesis^21,22,24,34–36^. These miRNAs are small non-coding RNAs that play a crucial role in shaping the hematopoietic landscape during developmental and adult hematopoiesis^37,38^. Thus, our hypothesis was that miRNAs carried by the AA BM-MSC EVs have a role in inhibiting the hematopoietic functions of HSPC in the BM niche. An altered expression of miRNAs in T-cell^39^ and plasma-derived exosomes^29^ has been reported in AA. However, studies on miRNAs in AA BM-MSC EVs is scarce. Using the NGS based miRNA profiling, the present study demonstrates that EVs from AA BM-MSC have a distinct miRNA profile compared to their normal counterparts. A total of 57 miRNAs, including precursor and mature miRNAs, were significantly deregulated in AA BM-MSC EVs. These results align with the previous studies, which have reported differential expression of miRNAs in the BM-MSC derived EVs from hematopoietic disorders such as AML^22^, MDS^23,36^, and multiple myeloma^24^. Additionally, the EV miRNA expression profile of our study was in alignment with the BM-MSC exosomal miRNA expression profile as reported by Ferguson et al., 2018^39^. However, it was completely different from the miRNA expression profile of plasma-derived exosomes as reported by Giudice et al.,2018^29^. Thus, highlighting that there could be a disparity in miRNA packaging in the EVs from different sources, signifying the presence/existence of a selective mechanisms for intercellular communication by EVs from different sources.

Since miRNAs from BM-MSC derived EVs impart their regulatory role by altering the genes and associated pathways at the post-transcriptional level, as described in hematological malignancies^24,35,36^. We tried to elucidate the functional role of these AA BM-MSC EVs encapsulated DE miRNAs by pathway enrichment analysis. To date two studies have been conducted to study the differential gene expression in HSPC in AA patients using a single cell genomics approach^40,41^. These studies demonstrated that HSPC from AA patients have upregulated genes involved in cell death, cytokine signalling, and immune responses and downregulated genes involved in cell cycle and cell differentiation. Further, the HSPC of AA patients had several changes linked to DNA damage and repair^41^. However, whether EV miRNAs impart these functional defects in HSPC is elusive in AA. Our GSEA analysis of significantly altered miRNAs showed enrichment of apoptosis and DNA damage response pathways. Additionally, pathways analysis showed enrichment of several pathways such as apoptosis, MAPK, PI3K/Akt, mTOR, chemokine receptor signalling, etc., which are crucial for maintaining hematopoietic homeostasis^42^. Our findings here provide with an insight that miRNAs enriched in AA BM-MSC EVs could potentially dampen HSPC functions in the microenvironment through impediment of these pathways, which is in alignment with our previous finding where we showed that AA BM-MSC EVs promoted apoptosis in healthy HSPC^25^. Concomitantly, our in-silico analysis of miRNA-mRNA interaction revealed that these EV miRNAs targeted 235 HSPC genes and their pathways analysis showed significant enrichment of pathways involved in cell survival and cell death. Altogether, these findings highlight the involvement of EV miRNAs of AA BM-MSC in the regulation of HSPC through cell survival and apoptotic mechanisms. However, the findings need to be established in different model systems, such as *in-vivo* and 3D scaffold, to study the in-depth interaction of these cells in the hematopoietic niche^43,44^.

To further comprehend the complex regulation and functional importance of HSPC genes in AA patients and role of BM-MSC EVs miRNA in their regulation, we performed hub gene analysis to identify the top 10 hub HSPC genes and their associated pathways. The functional analysis of Hub genes showed significant enrichment of pathways linked with cell cycle, survival, and cell death pathways. We made a Venn diagram to identify the pathways that were consistently enriched in dysregulated EV miRNAs, target HSPC and, hub genes. We found eight intersecting pathways: chemokine, MAPK, PI3K-Akt, Ras, mTOR, GABAergic synapse, cholinergic synapses and apoptotic signalling pathways targeted **Supplementary Fig.S5 and Table S4.** The role of the hub HSPC genes and their intervening pathways has been previously reported to be involved in regulating HSPC via different mechanisms as summarized in **Fig. S7** and **Supplementary Table S5** however, the role of these genes and associated miRNAs needs experimental validation in AA.

**Figure S7:**
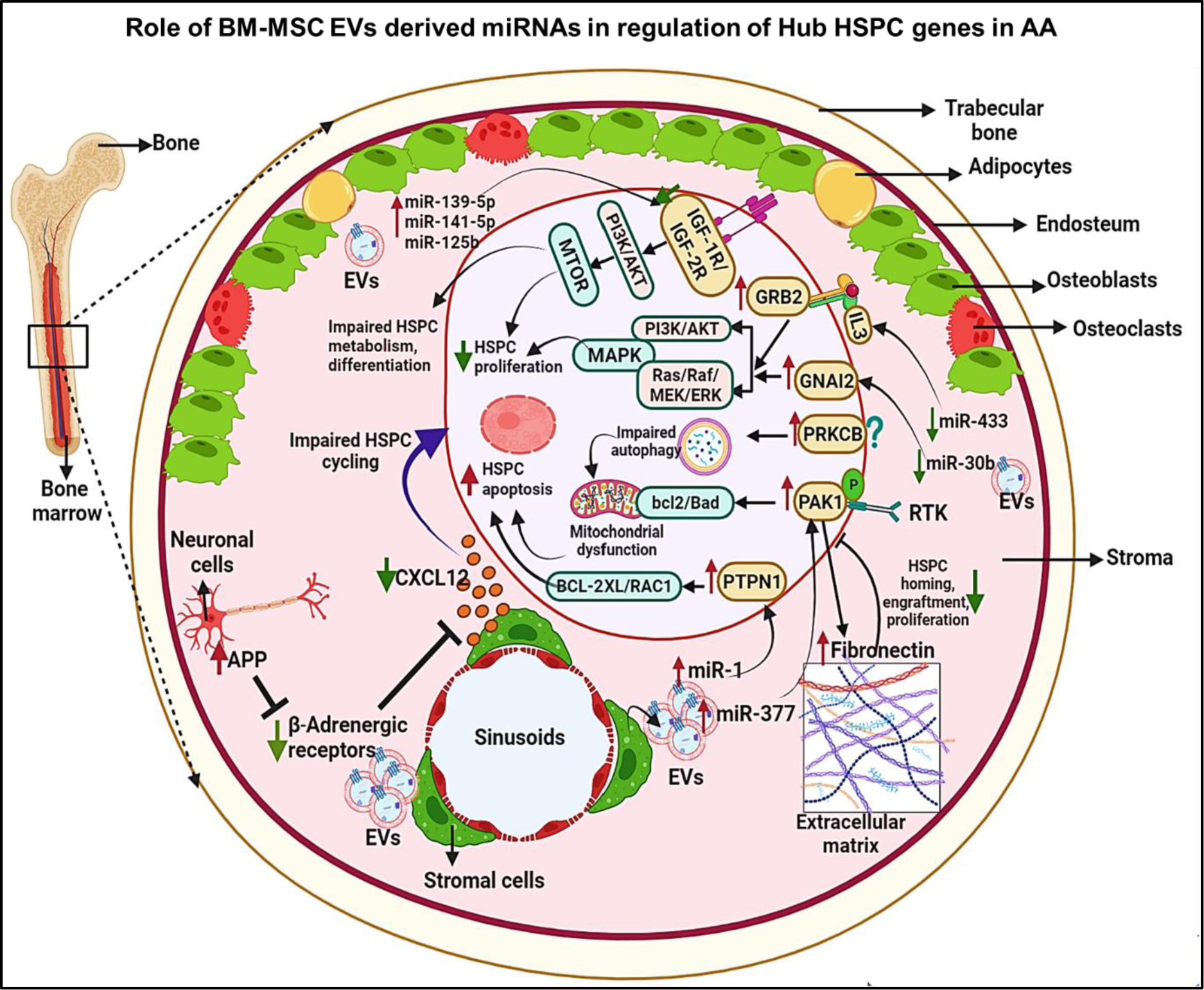
Diagrammatic illustration of cellular crosstalk mediated via AA BM-MSC EVs miRNAs in regulation of hub HSPC functions in the BM stroma of AA patients. The topmost hub-gene, APP is involved in the inactivation of β2-adrenergic receptor which could elicit a negative effect on HSPC migratory potential by supressing CXCL12. PAK1 activation is mediated via downregulation of miR-377 which further leads to the supressed fibronectin (an extracellular matrix protein) production. Moreover, depletion of fibronectin is associated with impaired HSPC homing, engraftment, proliferation, and differentiation. Furthermore, enhanced expression of PTPN1 can potentially reduce the HSPC number by activating apoptosis via suppression of RAC1 and BCL-XL. The role of PRKCB has not been studied in context of hematopoiesis however its role has been clearly demonstrated in inhibiting proliferation, promoting apoptosis, impaired mitochondrial dynamics and inhibition of autophagy in different diseases, these properties are hallmark of ageing of HSPC. Next, GNAI2 is a GPCR signalling protein that is modulated by RGS protein which is responsible for enhanced chemokine signalling, complement activation, migration and poor engraftment of HSPCs via PI3K/AKT and ERK/MAPK signalling. Moreover, GNAI2 has been identified as the direct target of miR-30b. Enhanced expression of Grb2 could induce a myeloid-bias of HSPC via IL3/ERK/MAPK signalling and have also been shown to be targeted by miR-433. Indispensable role of Insulin like Growth Factor-1 receptor (IGF-1R) and IGF-2R has been demonstrated in HSPC survival. In this study, decrease in IGF-1R and IGF-2R and upregulation of miR-125b, miR-141 and, miR-139-5p could be responsible for compromised HSPC function in AA. Overall, the study suggests a strong implication of miRNAs from BM-MSC EVs in deregulation of HSPC functions in AA via engagement of GABAergic synapse, cholinergic synapses, mTOR, MAPK pathways. Thus, our finding suggests that BM-MSC EV could potentially control the fate of HSPC via microRNA-mRNA regulatory axis.

Overall, in continuation of our previous study^25^, we now explicitly demonstrate that EVs from AA BM-MSC have differential regulation of miRNAs. We further demonstrate that these altered miRNAs can potentially target HSPC genes of AA patients and attenuate normal HSPC functions such as their proliferation, differentiation, and apoptosis by targeting several pathways. However, these miRNA-mRNA regulatory mechanisms need to be established in *in-vitro* and *in-vivo* settings to understand their role in AA pathobiology. Also, more studies are warranted on studying the functionality of BM-MSC derived EV-miRNA in a larger patient cohort and in the development of newer therapeutic strategies such as administration of engineered BM-MSC EVs in AA patients for the management of the disease. Nonetheless, our results also paved the way for exploring new signalling pathways such as GABAergic synapses, cholinergic synapses, and mTOR, which can be targeted for rescuing HSPC function in AA patients. Together, these findings provide valuable insight into the fact that the BM-MSC-derived EVs encapsulated miRNAs and their gene regulatory pathways are potentially involved in suppressing HSPC functions in AA. Thus, in the next step it would be important to investigate the pathways targeted by the differentially expressed miRNAs in the BM-MSC EVs of AA patients, which may be potential alternative therapeutic targets in AA.

## 4. Methodology

### 4.1. Study subjects

In the study six aplastic anemia (AA) patients and six age and gender-matched normal controls (n=6) were recruited. Diagnosis and severity of AA were determined by published criteria^45^. After informed consent, 5.0 ml of BM aspirate was collected in the Department of Hematology, Sanjay Gandhi Post graduate Institute of Medical Sciences, Lucknow. The procedures followed ethical standards of the Declaration of Helsinki 1975 and the study was approved by the Institutional Ethics Committee (IEC) and Institutional committee for Stem Cell Research (IC-SCR) of the Institute.

### 4.2. Isolation and characterization of Bone Marrow mesenchymal stromal cell (BM-MSC)

The bone marrow samples obtained from SAA patients and controls was processed using ficoll-density centrifugation method and mesenchymal stromal cells (MSCs) were cultured following standard laboratory protocol^11^. In brief, bone marrow mononuclear cells were cultured in the MSC supportive medium containing low glucose DMEM, 16% Fetal Bovine Serum-MSC Qualified (FBS), 1% Penicillin-Streptomycin and 1% Glutamax at 37°C under normoxic condition (All items were procured from Gibco, USA). Further, BM-MSC were characterized following the ISCT standards^46^. The cells of P3-P4 passage were used in the study for all the experiments.

### 4.3. Isolation of Extracellular vesicles (EVs) from BM-MSC

EVs were isolated from the BM-MSC using the Total Exosome Isolation (TEI) Reagent (Invitrogen, Life Technologies, Carlsbad, CA). When the BM-MSC in P3 passage reached a confluency of 80% the medium was replenished with exosome-free media and the cells were incubated in CO2 incubator at 37°C. After 48 hours to obtain EVs, cell culture supernatants were harvested and centrifuged at 2000 × g for 30 min to remove cells and debris. TEI reagent was added to the culture supernatant in a ratio of 1:2 in a new tube and was incubated overnight on a rotator mixer at 4 °C. Next day, the mixture was centrifuged at 10,000 × g for 1h at 4 °C and EV pellet was resuspended in in 1X phosphate buffered saline (PBS) and stored at -80°C for further experiments.

### 4.4. Nanoparticle Tracking Analysis (NTA)

For size distribution and particle concentration analysis of BM-MSC derived EVs, NTA was performed on Nanosight (NS300, Malvern Panalytical Ltd). In brief, EV preparation was diluted with 0.22µm pre-filtered 1X PBS in a ratio of 1:100. The diluted sample was then loaded onto the Nanosight chamber with a syringe pump. The motion of particles was acquired as three 60 second videos for each sample, and the data obtained was analyzed by NanoSight NTA 2.3 software for the estimation of modal particle size and concentration per ml.

### 4.5. Transmission Electron Microscopic (TEM) Analysis

A total of 10μl exosome suspensions were loaded on a carbon coated copper grid (Sigma-Aldrich, USA) and desiccated for 20 min. Then, 2% uranyl acetate in water was added to the exosome layer and was allowed to dry overnight. After that, TEM imaging was performed the next day on TEM-1400 Plus, JOEL (JEOL, Japan).

### 4.6. Western Blotting

Exosomes were lysed by RIPA buffer (BioRad, USA) and the protein concentrations were obtained by Pierce BCA Protein Assay (Thermo Scientific, USA). Extracted proteins were separated in 12% polyacrylamide gel electrophoresis and analyzed by Western blot. Proteins were transferred to PVDF membranes and were incubated with CD63 (1:1,000, Abcam), CD81 (1:1,000, Abcam), TSG101 (1:1,000, Abcam) and Calnexin (1:1,000, Santa Cruz) at 4°C overnight (Details of antibodies are provided in supplementary data) overnight at 4 °C. Next day, membrane was incubated with (Horseradish peroxide) HRP labelled secondary antibody following washing with secondary antibody, the proteins bands were visualized by chemiluminescence (Invitrogen, USA).

### 4.7. Total RNA isolation, microRNA profiling and, Data analysis

RNA was isolated from the BM-MSC EVs using total Exosome RNA and Protein Isolation Kit (Invitrogen) following manufacturer’s instruction. The RNA concentration was analysed using Nanodrop Spectrophotometer 2000 (Thermo-fisher Scientific). The sample were analyzed for their integrity based on RIN number as analysed on bioanalyzer. After qualifying Quality QC samples were processed for library preparation and QC. The raw data fastq sequences were generated and demultiplexed using CASAVAv1.8 pipeline. We implemented a comprehensive RNA-Seq analysis pipeline utilizing the STAR v2.7.10a (Spliced Transcripts Alignment to a Reference) method for the alignment of sequencing reads^47^. The quality control of raw reads was assessed by checking md5sum files, and further evaluation was conducted using FastQC v0.11.7 to ensure data integrity. Adapter trimming was executed with the fastp v0.23.2 to enhance data quality^48^. The STAR method was employed for aligning reads to human reference hg38 using GTF file “gencode.v42.basic.annotation.miRNA.gtf”, incorporating both raw and mature sequences derived from has.gff files sourced from the miRDB^49^. Subsequently, the filtered reads were aligned against this constructed miRNA genome. The quantification of individual miRNA expression levels was achieved through the utilization of the feature counts v2.0.3, providing a robust and reliable assessment of miRNA expression profiles in the context of our RNA-Seq analysis^50^. This pipeline not only ensures the accuracy and reliability of the obtained results but also establishes a standardized approach for the investigation of miRNA expression in our experimental system. The raw data files for 10 samples have been submitted in NCBI under BioProject number PRJNA1094675 (SRA accession no: SRR28513789, SRR28513791, SRR28513790, SRR28513792, SRR28513788, SRR28513785, SRR28513787, SRR28513784, SRR28513786, SRR28513783)

### 4.8. Differential expression of BM-MSC EV derived microRNA and functional analysis

To identify the differentially expressed miRNAs in the EVs of AA BM-MSC in comparison to control the DESeq2 v1.43.1tool of the R Bioconductor package was used^51^. The differentially expressed miRNAs were selected based on *p-value* <0.05 and the absolute value of log2fold change (log2FC) >1.0 (up-regulated) or < -1.0 (down-regulated)^44,52,53^. Expression profile of deregulated miRNAs was represented in the heatmap using Complex Heatmap tool (2.12.1version) integrated in R bioconductor.

### 4.9. Data processing and differential expression analysis of GEO dataset

To identify the potential HSPC target genes of differentially expressed miRNAs we analysed GSE165870^41^ dataset by DEseq2 as mentioned above. To identify significant DE genes, we applied the threshold of *adjusted p-value* <0.05 and log2FC > 1.0 (upregulated) or log2FC < −1.0 (downregulated).

### 4.10. Functional enrichment analysis and Target gene identification of differentially expressed EV miRNAs

For functional enrichment analysis of differentially expressed miRNAs, we used the miRNA Enrichment Analysis and Annotation Tool (miEAA), a web-based tool that facilitates the functional analysis of sets of miRNAs^31^. The tool has integrated Gene ontology (GO), wikipath, Reactome and KEGG databases for GO and pathway analysis. The GO and pathways having *p-value* <0.05 were considered statistically significant. The experimentally validated target genes of DE-miRNAs were predicted using miRNet 2.0, which is a comprehensive tool that integrates data from eleven different miRNA databases (TarBase, miRTarBase, miRecords, miRanda, miR2Disease, HMDD, PhenomiR, SM2miR, PharmacomiR, EpimiR and starBase)^32^. We also used miEAA for the Gene Set Enrichment Analysis (GSEA)^54,55^ of the DE miRNAs.

### 4.11. Identification of candidate target genes and miRNA-mRNA Regulatory Network Construction

To identify the candidate target genes, the common target identified from different miR databases and differentially expressed HSPC genes (GSE165870). With the help of the miRNet 2.0^32^, a miRNA-mRNA regulatory network was established and visualized.

The tool provides the graphical visualization of the interaction between dysregulated EV miRNAs and mRNAs.

### 4.12. Protein-Protein Interaction Network Construction and Hubb gene identification

The Protein-protein interaction (PPI) of differentially expressed target genes was performed using online STRING^56^, which predicts proteins interaction and associated pathways. A combined score of >0.7 was set as threshold for PPI network construction and visualization by Cytoscape. Another plug-in of Cytoscape, molecular complex detection (MCODE) was employed to identify key gene modules^57,58^. Further, the top ten hub genes were identified and visualized using cyto-hubba plugin of Cytoscape. The size of the node is represented by the degree value. The GO and KEGG pathway analysis of hub genes was performed using DAVID.

### 4.13. Data visualisation

A volcano map was executed in R software for the miRNA and mRNA data with the ggplot2 package in R. DE miRNA expression profile was represented in the heatmap using ComplexHeatmap package. Chord plot was used to represent the interaction between miRNAs and pathways, and gene and pathways. Sankey plot was used to represent DE mRNA, enriched pathways, and their gene ratio. Alluvial plot was used to represent miRNAs and pathways which are involved in regulation of hub genes. For data visualisation R Bioconductor and SR plots (https://www.bioinformatics.com.cn/srplot) was used to enhance the representation of the data.

## Acknowledgement

We would like to thank Dr. Khaliqur Rehman, Additional Professor, Dept. of Hematology, SGPGIMS, for helping with the flow cytometry experiments, and Mr. Ali and Mrs. Neetika Mishra, lab-technicians, Dept. of Hematology, SGPGIMS for their help in sample collection and processing. We would also like to express our deepest gratitude to all patients who consented to participate in this study.

## 7. Authors contribution

J.S.–Conceived and designed the project, performed wet lab experiments, data analysis and wrote the manuscript; K.K.-performed the seq analysis and proof-read the bioinformatic part in the manuscript under the supervision of R.K.; B.R. and P.S.-performed with in-vitro experiments, in-silico analysis of data; N.T., R.G., and S.Y.-helped with the collection of patient sample along with their diagnostic evaluation and, S.N. and C.P.C. Conceived, designed and initiated the project, provided guidance during data analysis, proofread the whole manuscript, and supervised the project. All authors read and approved the final version of the manuscript.

## 8. Data availability

The raw data files for 10 samples have been submitted in NCBI under BioProject number PRJNA1094675. R scripts for the analysis of exosomal miRNA profiling and RNA-Seq transcriptomics data are available for downloading using the following link http://github.com/CGnTLab/miR.git. Any additional information can be availed from the corresponding authors upon reasonable request.

## 11. Ethical Approval

This study was conducted in accordance with the Declaration of Helsinki (as revised in 2013) and approved by the Institutional Ethics Committee (IEC-2018-173-EMP-107) and the Institutional Committee for Stem Cell Research (ICSCR–2019-02-EMP-08) of the Sanjay Gandhi Post Graduate Institute of Medical Sciences, Lucknow.

