## Supplementary Material for "Altered Expression of microRNAs Implicated in Hematopoietic Dysfunction in the Extracellular Vesicles of Bone Marrow-Mesenchymal Stromal Cells in Aplastic Anemia"

List of supplementary figures and tables:

| S.No. | Figure/Table No. | Description |
|---|---|---|

| | | |
|---|---|---|
| 1 | **Figure S1** | Immunophenotypic and differentiation characterization of AA and normal control (NC) BM-MSC. |
| 2 | **Figure S2** | TEM image showing particle size and original immunoblots. |
| 3 | **Figure S3** | Differentially expressed EV miRNAs and HSPCs genes regulatory network analysis performed using miRNet 2.0 tool. |
| 4 | **Figure S4** | Gene ontology analysis of Hub genes. |
| 5 | **Figure S5** | Intersecting pathways between differentially expressed miRNA, Target HSPCs, and Hub genes. |
| 7 | **Table S1** | Differentially expressed precursor miRNA in EV from AA BM-MSC with fold change and p-values. |
| 8 | **Table S2** | Differentially expressed mature miRNA in EV from AA BM-MSC with fold change and p-values. |
| 10 | **Tabe S3** | Details of GO cellular component and molecular functions of the hub genes. |
| 11 | **Table S4** | Direction of regulation or Log2FC expression of Hub genes and their associated AA BM-MSC EVs miRNAs. |
| 12 | **Table S5** | Role of hub genes and associated pathways in regulation of HSPC functions. |

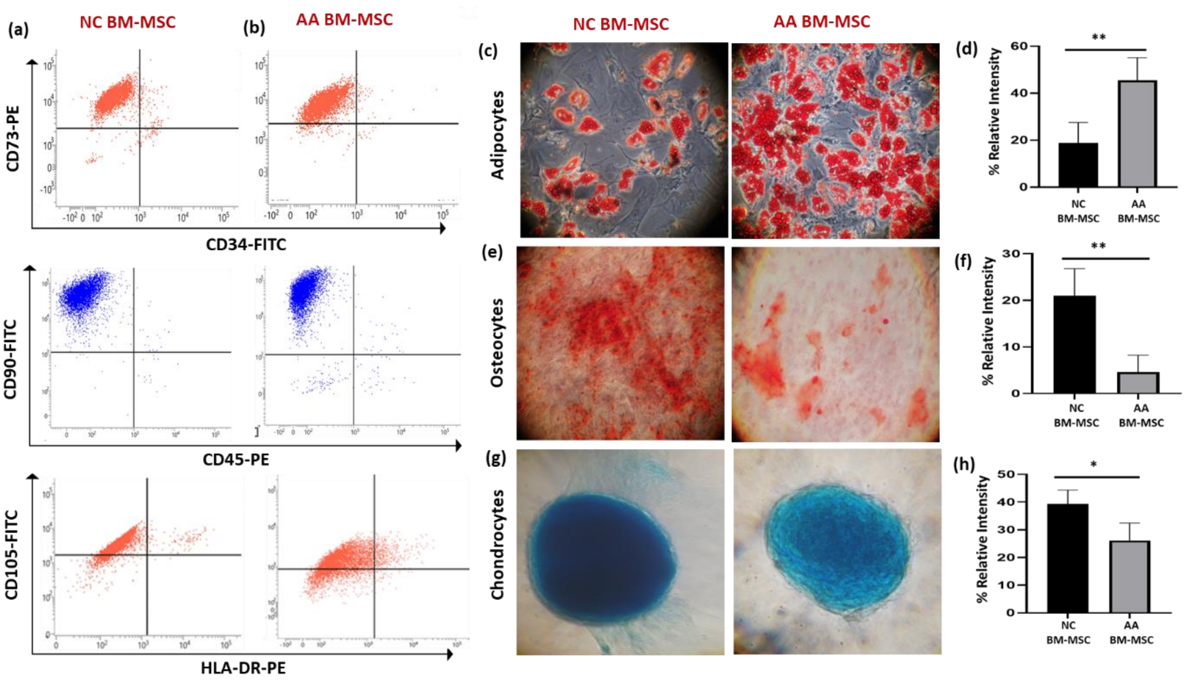


**Figure S1**: **Immunophenotypic and differentiation characterization of AA and normal control (NC) BM-MSC**. Representative flow cytometry dot plots of, **(a)** AA BM-MSC and **(b)** NC BM-MSC showing presence of CD73, CD90 and CD105 and no expression of CD34, CD45 and HLA-DR. Representative photomicrograph and respective bar graphs exhibiting the adipogenic differentiation **(c-d),** osteocytic differentiation potential **(e-f)** and, chondrogenic differentiation potential **(g-h)** in AA and control BM-MSC. Note: CD-cluster of differentiation; *p value*: <0.05*; <0.01**; and <0.001***

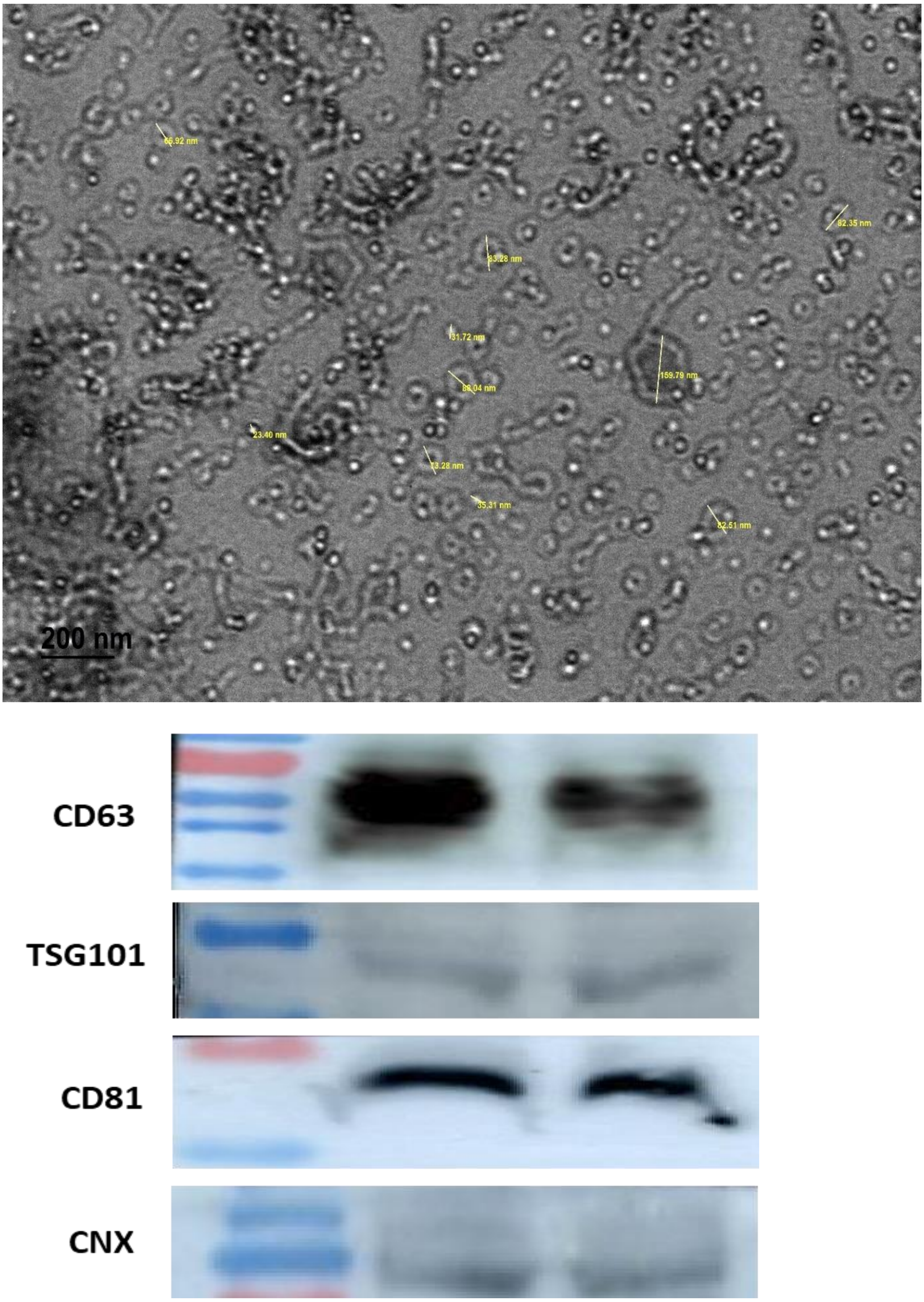


**Figure S2: TEM image showing particle size and original immunoblots.** Characterization of AA and control BM-MSC EVs using western blot. Original western blot images of CD63, TSG101, CD81 and Calnexin (CNX)

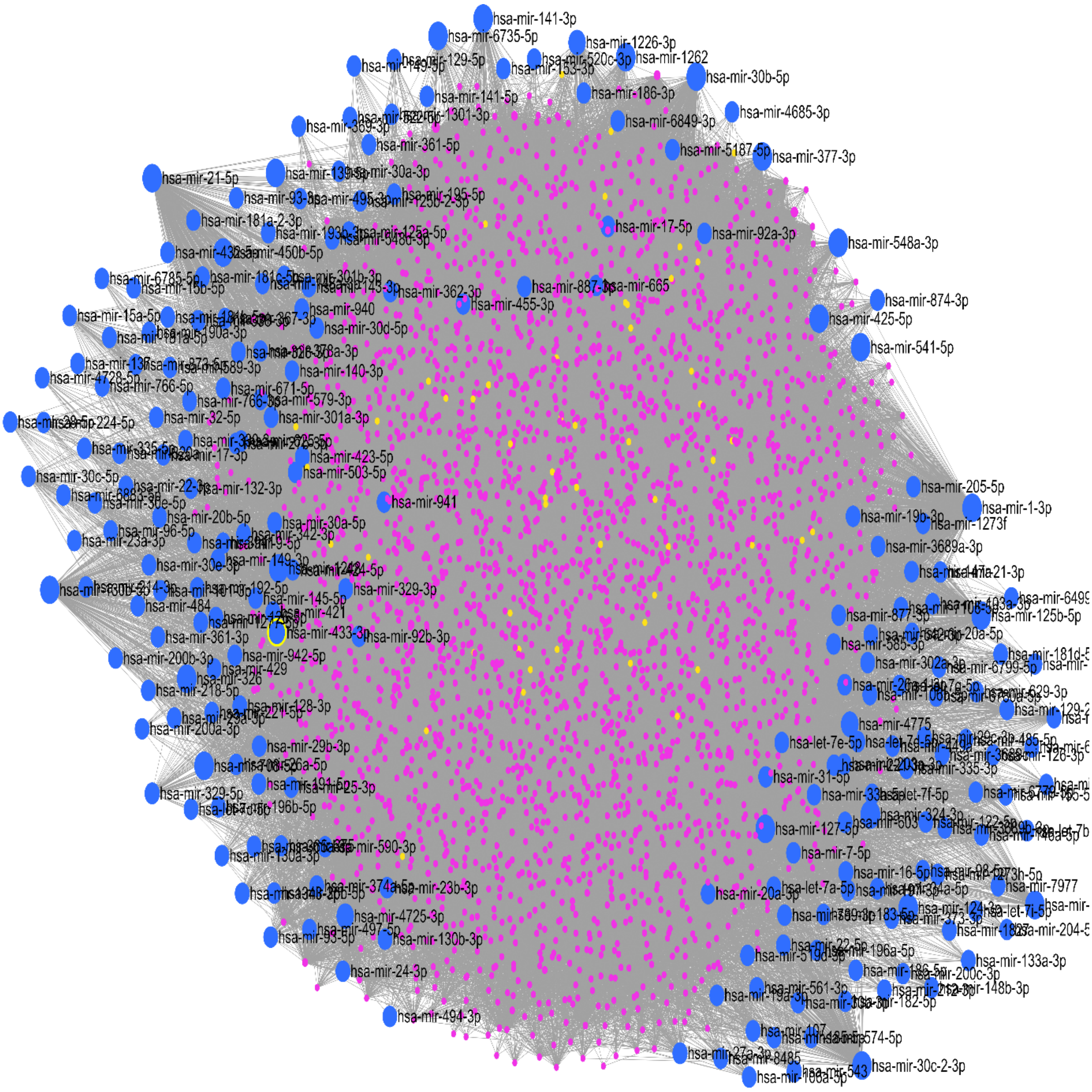


**Figure S3: Differentially expressed EV miRNAs and HSPCs genes regulatory network analysis performed using miRNet 2.0 tool.** The blue dots represent the seed miRNAs and interacting miRNAs and the pink dots are seed mRNA and interacting mRNAs.

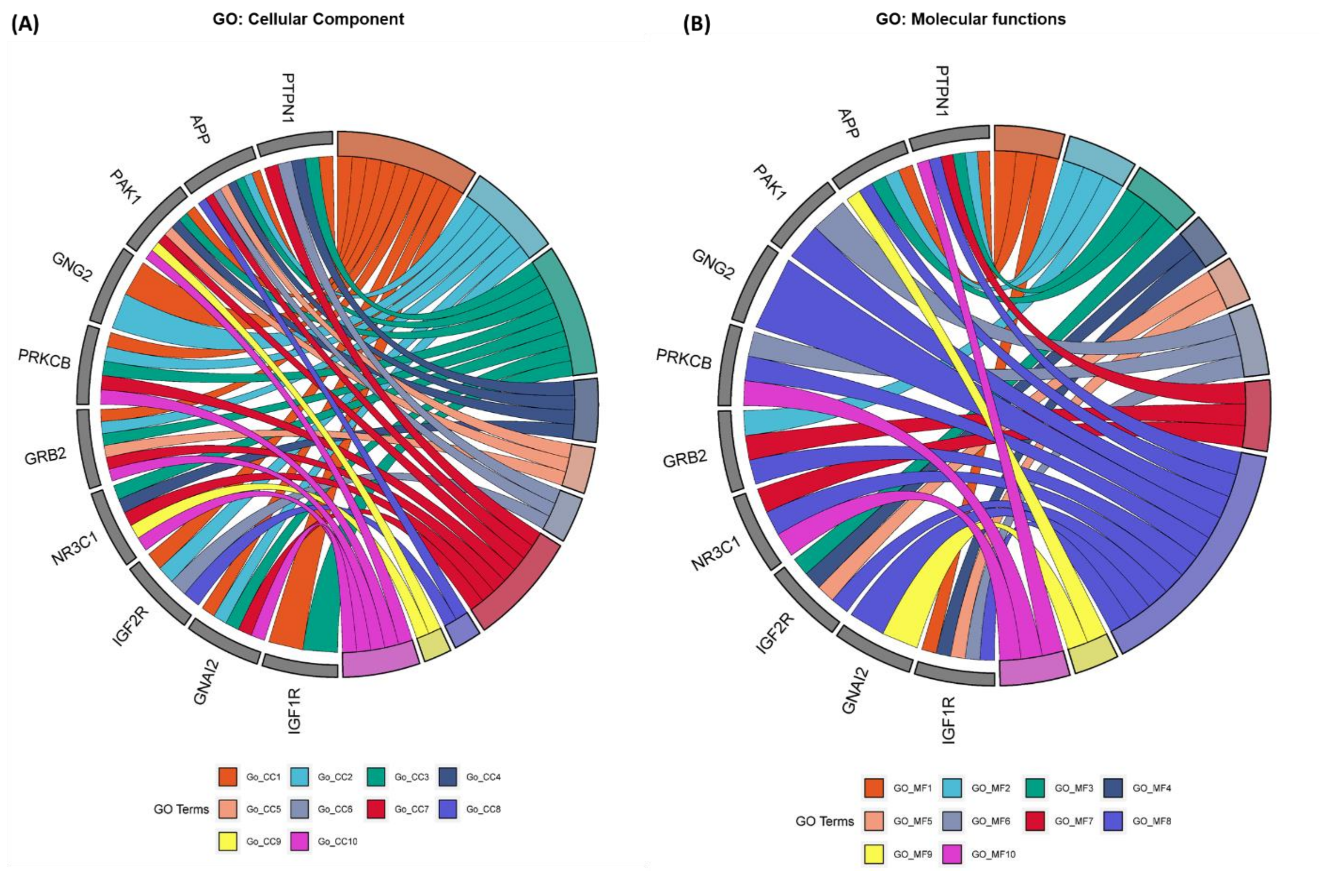


**Figure S4: Gene ontology analysis of Hub genes. (A)** Top 10 GO cellular component of the Hub genes and, **(B)** Top 10 GO molecular functions of the Hub genes. **Note:** The detail of GO cellular component and molecular functions and mentioned in the figure legend is mentioned in the Table S4.

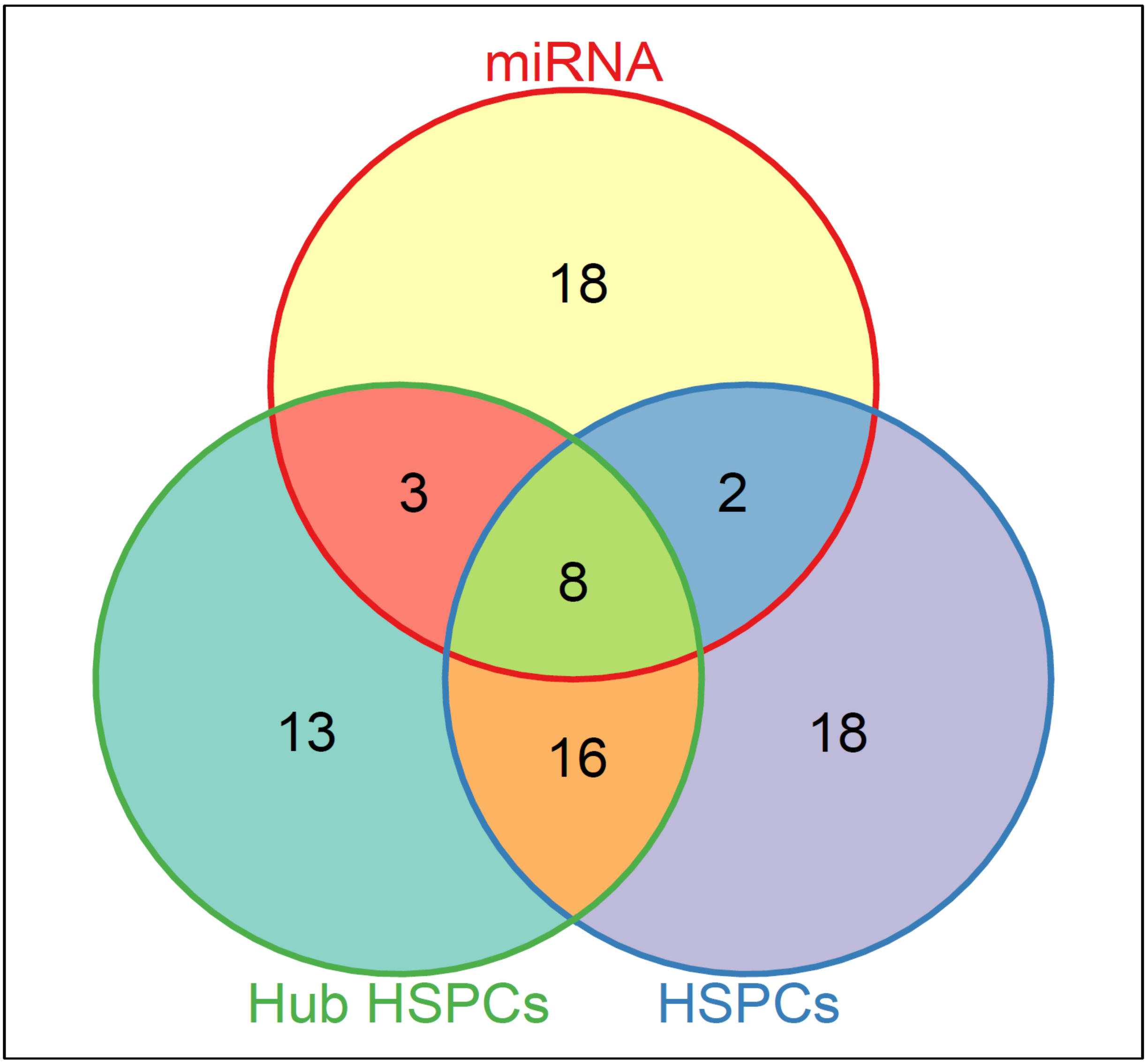


**Figure S5: Intersecting pathways between DE miRNA, Target HSPCs, and Hub genes.** The representative Venn diagram showing the intersecting pathways, the 8 pathways that were common in all the three groups were viz., Chemokine, MAPK, PI3K-Akt, Ras, mTOR, GABAergic synapse, cholinergic synapses and, apoptotic signalling pathways.

**Table S1:** Differentially expressed precursor miRNA in EV from AA BM-MSC with fold change and p-values.

| miRNA | log2 Fold Change | p-value |
|---|---|---|
| hsa-mir-326 | 23.73942086 | 8.88E-15 |
| hsa-mir-4775 | -21.55684553 | 4.75E-12 |
| hsa-mir-6735 | -20.94080044 | 1.89E-11 |
| hsa-mir-141 | 8.858673661 | 0.001476 |
| hsa-mir-30b | -9.033726703 | 0.001413 |
| hsa-mir-1226 | -8.054840038 | 0.002588 |
| hsa-mir-874 | -7.984082846 | 0.002526 |
| hsa-mir-433 | -7.98934831 | 0.003232 |
| hsa-mir-708 | -8.303398288 | 0.006259 |
| hsa-mir-4725 | 7.958170327 | 0.009382 |
| hsa-mir-10395 | 2.648795643 | 0.013188 |
| hsa-mir-21 | -1.246678951 | 0.017018 |
| hsa-mir-425 | -1.268264206 | 0.016929 |
| hsa-mir-450b | -7.298357251 | 0.019243 |
| hsa-mir-887 | -7.158619707 | 0.021683 |
| hsa-mir-1262 | -6.817874644 | 0.028793 |
| hsa-mir-585 | -6.646522572 | 0.033073 |
| hsa-mir-125b-1 | 0.946374758 | 0.036104 |
| hsa-mir-125b-2 | 0.949233181 | 0.03551 |
| hsa-mir-5187 | -6.401988058 | 0.040118 |
| hsa-mir-541 | -6.338096222 | 0.042156 |
| hsa-mir-139 | 6.094656352 | 0.044456 |

**Table S2:** Differentially expressed mature miRNA in EV from AA BM-MSC with fold change and p-values.

| miRNA | log2 Fold Change | p-value |
|---|---|---|
| hsa-miR-326 | 23.73942086 | 8.88E-15 |
| hsa-miR-543 | -22.6464159 | 3.74E-13 |
| hsa-miR-26a-1-3p | -21.7728567 | 2.89E-12 |
| hsa-miR-377-3p | -21.53978926 | 4.94E-12 |
| hsa-miR-4775 | -21.55684553 | 4.75E-12 |
| hsa-miR-6735-5p | -20.94080044 | 1.89E-11 |
| hsa-miR-141-3p | 8.858673661 | 0.001476 |
| hsa-miR-30b-5p | -9.033726703 | 0.001413 |
| hsa-miR-130b-5p | -7.748026227 | 0.00176 |
| hsa-miR-425-5p | -1.584965272 | 0.002656 |
| hsa-miR-874-3p | -7.951251139 | 0.002772 |
| hsa-miR-433-3p | -7.98934831 | 0.003232 |
| hsa-miR-6716-3p | 8.120172313 | 0.008026 |
| hsa-miR-708-5p | -8.163320974 | 0.008286 |
| hsa-miR-4725-3p | 7.958170327 | 0.009382 |
| hsa-miR-1-3p | 7.851507909 | 0.010383 |
| hsa-miR-324-3p | -7.975105575 | 0.01052 |
| hsa-miR-329-5p | -7.372111851 | 0.013757 |
| hsa-miR-138-5p | -7.007447463 | 0.018015 |
| hsa-miR-450b-5p | -7.298357251 | 0.019243 |
| hsa-miR-548a-3p | -7.362080411 | 0.018212 |
| hsa-miR-10395-3p | 2.422679292 | 0.022213 |
| hsa-miR-10395-5p | 2.604258124 | 0.022776 |
| hsa-miR-1226-3p | -7.208892555 | 0.020776 |
| hsa-miR-21-5p | -1.272369076 | 0.023025 |
| hsa-miR-887-3p | -7.158619707 | 0.021683 |
| hsa-miR-1248 | 4.656750356 | 0.028449 |
| hsa-miR-1262 | -6.817874644 | 0.028793 |
| hsa-miR-585-3p | -6.646522572 | 0.033073 |
| hsa-miR-125b-5p | 0.9543204 | 0.035228 |
| hsa-miR-30c-2-3p | -6.392281797 | 0.040422 |
| hsa-miR-5187-5p | -6.401988058 | 0.040118 |
| hsa-miR-541-5p | -6.338096222 | 0.042156 |
| hsa-miR-127-5p | -6.259487546 | 0.044784 |
| hsa-miR-139-5p | 5.297119226 | 0.04917 |

**Tabe S3:** Details of GO cellular component and molecular functions of the hub genes.

| GO TERM | Term | Count | P-Value | Genes | FDR |
|---|---|---|---|---|---|
| Go_CC1 | GO:0005886~plasma membrane | 9 | 1.38E-04 | PTPN1, APP, PAK1, GNG2, PRKCB, GRB2, IGF2R, GNAI2, IGF1R | 0.012701 |
| Go_CC2 | GO:0070062~extracellular exosome | 6 | 0.001246 | APP, GNG2, PRKCB, GRB2, IGF2R, GNAI2 | 0.054149 |
| Go_CC3 | GO:0005737~cytoplasm | 8 | 0.002511 | PTPN1, APP, PAK1, PRKCB, GRB2, NR3C1, GNAI2, IGF1R | 0.054149 |
| Go_CC4 | GO:0032991~macromolecular complex | 4 | 0.002638 | PTPN1, APP, PAK1, NR3C1 | 0.054149 |
| Go_CC5 | GO:0005911~cell-cell junction | 3 | 0.002943 | APP, PAK1, GRB2 | 0.054149 |
| Go_CC6 | GO:0005769~early endosome | 3 | 0.007608 | PTPN1, APP, IGF2R | 0.087205 |
| Go_CC7 | GO:0005829~cytosol | 7 | 0.015178 | PTPN1, APP, PAK1, PRKCB, GRB2, NR3C1, GNAI2 | 0.102538 |
| Go_CC8 | GO:0032588~trans-Golgi network membrane | 2 | 0.044488 | APP, IGF2R | 0.255804 |
| Go_CC9 | GO:0005815~microtubule organizing center | 2 | 0.047706 | PAK1, NR3C1 | 0.366411 |
| Go_CC10 | GO:0005654~nucleoplasm | 5 | 0.049708 | PAK1, PRKCB, GRB2, NR3C1, GNAI2 | 0.407398 |
| GO_MF1 | GO:0005158~insulin receptor binding | 3 | 5.05E-05 | PTPN1, APP, IGF1R | 0.004631 |
| GO_MF2 | GO:0046875~ephrin receptor binding | 3 | 9.26E-05 | PTPN1, APP, GRB2 | 0.004631 |
| GO_MF3 | GO:0019899~enzyme binding | 2 | 0.01316 | IGF2R, IGF1R | 0.123203 |
| GO_MF4 | GO:0005520~insulin-like growth factor binding | 2 | 0.010876 | IGF2R, IGF1R | 0.164497 |

| | | | | | |
|---|---|---|---|---|---|
| GO_MF5 | GO:0001965~G-protein alpha-subunit binding | 3 | 0.012756 | PTPN1, APP, IGF2R | 0.164497 |
| GO_MF6 | GO:0004712~protein serine/threonine/tyrosine kinase activity | 3 | 0.01739 | PAK1, PRKCB, IGF1R | 0.193223 |
| GO_MF7 | GO:0019901~protein kinase binding | 3 | 0.023993 | PTPN1, GRB2, NR3C1 | 0.239299 |
| GO_MF8 | GO:0005515~protein binding | 10 | 0.026323 | PTPN1, APP, PAK1, GNG2, PRKCB, GRB2, NR3C1, IGF2R, GNAI2, IGF1R | 0.239299 |
| GO_MF9 | GO:0001664~G-protein coupled receptor binding | 2 | 0.033235 | APP, GNAI2 | 0.276957 |
| GO_MF10 | GO:0008270~zinc ion binding | 3 | 0.041569 | PTPN1, PRKCB, NR3C1 | 0.473606 |

**Table S4:** Direction of regulation or Log2FC expression of Hub genes and their associated AA BM-MSC EVs miRNAs.

| S. No. | Hub Genes | Log2FC expression of Hub genes | miRNAs | Log2FC expression of miRNAs |
|---|---|---|---|---|
| **1** | APP | 1.69 | miR-548a | -7.36 |
| **2** | GRB2 | 1.22 | miR-324 | -7.98 |
| | | | miR-433 | -7.99 |
| | | | miR-541 | -6.31 |
| **3** | GNG2 | 1.22 | miR-138-5p | -7.01 |
| **4** | GNAI2 | | miR-138-5p | -7.01 |
| | | | miR-30b-5p | -9.03 |
| | | | miR-1-3p | 7.85 |
| **5** | PAK1 | 1.14 | miR-377-3p | -21.54 |
| **6** | PTPN1 | 1.78 | miR-1-3p | 7.85 |
| **7** | NR3C1 | 1.04 | miR-377-3p | -21.54 |
| | | | miR-127-5p | -6.26 |
| **8** | PRKCB | 0.89 | miR-433-3p | -7.99 |
| | | | miR-130b-5p | -7.75 |
| | | | miR-326 | 23.73 |
| **9** | IGF1R | -1.96 | miR-139-5p | 5.29 |
| | | | miR-377-3p | -21.54 |
| | | | miR-125b-5p | 0.95 |
| | | | miR-141-3p | 8.85 |
| | | | miR-10395-3p | 2.42 |
| | | | miR-1226 | -7.21 |
| **10** | IGF2R | -3.03 | miR-377-3p | -21.54 |

**Table S5:** Role of hub genes and associated pathways in regulation of HSPC functions.

| S.No | Hub-Gene | Function | Reference |
|---|---|---|---|
| 1 | APP (Amyloid precursor protein) | • Highly reported in Alzheimer's disease, where it is involved in the inactivation of β2-adrenergic receptor.<br>• Allogenic transplantation of HSPC have been shown to rescue β2-adrenergic defect and APP expression.<br>• β2-adrenergic agonists have been shown to promote HSPC mobilization by stimulating G-CSF | 1,2,3,4 |
| 2 | GNAI2 (Guanine Nucleotide Binding Protein (G Protein), Alpha Inhibiting Activity Polypeptide 2) | • GPCR signalling protein that is modulated by RGS protein which is responsible for enhanced chemokine signalling, complement activation and poor engraftment of HSPC<br>• It's down-expression reduces hepatocellular carcinoma cell migration potential when targeted by miR-30b. | 5,6 |
| 3 | PAK1 (p-21 activated kinases) | • A serine threonine kinases that regulate several functions such as cytoskeletal remodelling, cell proliferation and apoptosis and mitosis[53]<br>• In diabetic nephropathy, activation of PAK1 by miR-377 leads to production of fibronectin.<br>• Fibronectin is an essential component of extracellular matrix and along with its receptor, it role has been reported to be associated with HSPC homing, engraftment, proliferation, and differentiation[56] | 7,8,9,10 |
| 4 | PTPN1 (Protein Tyrosine Phosphatase | • A member of protein tyrosine phosphatase family | 11 |

| | | | |
|---|---|---|---|
| | Non-Receptor Type 1) | • Depletion of PTPN1 is reported to improve HSPC number in BM and spleen significantly.<br>• Supresses HSPC apoptosis via activation of the RhoGTPase, RAC1, and induction of BCL-XL | |
| 5 | NR3C1 (Nuclear Receptor Subfamily 3 Group C Member 1) | • A Glucocorticoid Receptor involved in a complex interplay with G-CSF mediated via cholinergic signalling and regulates HSPC mobilization | 12 |
| 6 | GNG2 (Guanine Nucleotide Binding Protein Gamma 2) | • Not reported in context of regulating HSPC function but have role in cell proliferation.<br>• Reported to suppress proliferation of breast cancer cells via MRAS signalling. | 13 |
| 7 | PRKCB (Protein Kinase C Beta) | • Not reported in context of regulating HSPC function but have role in autophagy and mitochondrial dynamics.<br>• Activation of PRKCB negatively modulates the mitochondrial energy status and inhibits autophagy.<br>• Pharmacological inhibition of PRKCB improved autophagy via increase in the mitochondrial membrane potential. | 14 |
| 8 | Insulin-like Growth Factor-1 receptor (IGF-1R) | • Have indispensable role in HSPC survival<br>• Decline in IGF1 limits HSPC survival and hematopoietic health span.<br>• It is targeted by several miRNA miR-125b, miR-141, miR-139-5p and, miR-1226, miR-377 in different diseases. However miR-IGFI regulatory axis remains unexplored in AA. | 15,16,17,18,19,20 |
| 9 | Insulin-like Growth Factor-2 | • IGF2) to be predominantly expressed within long-term HSC | 21,22 |

| | | | |
|---|---|---|---|
| | receptor (IGF-2R) | • Preserves long-term HSC through upregulation of the CDKi, p57 via activation of the PI3K-Akt pathway | |
| 10 | GRB2 (Growth factor receptor-bound protein 2 ) | • Preferentially expressed in HSC compared to its progenitor cells.<br>• Deletion of Grb2 induced a rapid decline of erythroid and myeloid progenitors and a progressive decline of HSC numbers via IL3/ERK.MAPK signalling. | 23,24 |
